# Severe enamel defects in wild Japanese macaques

**DOI:** 10.1101/2023.05.18.541087

**Authors:** Ian Towle, Carolina Loch, Marina Martínez de Pinillos, Mario Modesto-Mata, Leslea J. Hlusko

## Abstract

Plane-form enamel hypoplasia (PFEH) is a severe dental defect in which large areas of the crown are devoid of enamel. This condition is rare in humans and rarer in wild primates. The etiology of PFEH has been linked to exposure to severe disease, malnutrition, environmental toxins, and other systemic conditions. Similar defects have also been associated with genetic conditions such as amelogenesis imperfecta. In this study, we examined the prevalence of all types of enamel hypoplasia in several populations of wild Japanese macaques (*Macaca fuscata*) with the aim of providing context for the severe defects found in macaques from Yakushima Island. We found that macaques from other islands and from the mainland of Japan have low prevalence of the more common types of enamel hypoplasia and none exhibit PFEH. In contrast, 10 of the 21 individuals (48%) from Yakushima Island displayed uniform and significant PFEH, with all 10 living in two adjacent locations in the south of the island. All permanent teeth were affected to varying degrees except for first molars, and the mineral content of the remaining enamel in teeth with PFEH is normal (i.e., no hypo- or hyper mineralization). Given that the affected individuals have smaller first molars compared to non-affected macaques, and that they all underwent dental development during a period of substantial human-related habitat change, we conclude that the PFEH likely resulted from environmental stress. Extreme weather events on the island may also have influenced the formation of these defects. Additionally, it is plausible that a documented recent population bottleneck could have heightened the susceptibility of these macaques to PFEH. Further research on living primate populations is needed to better understand the causes of PFEH in wild primates and to evaluate whether such features can be used to assess the impact of human-related disturbance.

## Introduction

Enamel hypoplasia is a common type of dental defect in which there is reduction in enamel volume caused by cessation or diminution of ameloblast function (Hillson and Bond, 1997; Xing et al., 2016; McGrath et al., 2018). Defects are often characterized into one of four categories: linear-form (LEH), pit-form (PEH), plane-form (PFEH), and localized enamel hypoplasia (Seow, 1990; Hillson and Bond, 1997; Guatelli-Steinberg, 2015; Skinner et al., 2016; Towle and Irish, 2020). The classification of defects into these categories can sometimes be difficult (e.g., Ogden et al., 2007; Ioannou et al., 2016; Towle et al., 2018; Towle and Irish, 2020). Enamel hypoplasia has been commonly recorded and described in archaeological and paleontological studies, with the ‘health’ or ‘stress’ of populations or taxa often assessed using hypoplasia as a proxy (e.g., Guatelli-Steinberg et al., 2004; Xing et al., 2016; Orellana-González et al., 2020; Minozzi et al., 2020). The use of enamel hypoplasia as a tool to understand disturbances to recent or contemporary populations has received much less attention (e.g., Brown et al., 2020), and its potential application to help assess habitat and environmental impacts remains relatively unexplored. Here, we investigate a particularly severe type of enamel hypoplasia and explore the information it may provide about a population of macaques born during a period of substantial human-induced habitat change on the island of Yakushima, Japan.

PFEH is one of the most severe and rarest types of enamel hypoplasia, resulting from a complete cessation of enamel matrix formation that results in large areas of the tooth crown having little or no enamel deposition (Hillson and Bond, 1997; Ogden et al., 2007; Sawada et al., 2008; Krenz-Niedbała and Kozłowski, 2013; Towle et al., 2017; Towle and Irish, 2020). Severe dental defects such as PFEH are rarely described in wild primates (Towle et al., 2018). Severe tooth defects in which large areas of enamel do not form correctly often relate to human-specific conditions, diseases and environmental factors such as congenital syphilis, medical treatments associated with the use of mercury, and specific genetic conditions such as amelogenesis imperfecta (Ogden et al., 2007; Crawford et al., 2007; Ioannou et al., 2016; Towle et al., 2017; Ioannou et al., 2018). These factors may have contributed to PFEH being more frequently described in humans compared to other primate taxa.

Less severe enamel defects, such as localized enamel hypoplasia, linear enamel hypoplasia (LEH), and pit-form hypoplasia (PEH) are, in contrast, relatively common in wild primates (e.g., Guatelli-Steinberg and Skinner, 2000; Hannibal and Guatelli-Steinberg, 2005; Newell et al., 2006; McGrath et al., 2018; Towle and Irish, 2020). The prevalence and severity of these defects varies between primate groups; not just based on illness, malnutrition, and disease susceptibility, but also relating to tooth developmental timing and morphological/structural differences among teeth (McGrath et al., 2018; Towle and Irish, 2020). Therefore, it can be difficult to macroscopically compare different taxa based on prevalence and severity of PEH, LEH and localized defects. Unlike these other types of enamel hypoplasia, in all cases PFEH are considered extreme defects, and are thought to be associated with life-threatening periods of malnutrition or disease (Ogden et al., 2007; Crawford et al., 2007; Ioannou et al., 2016; Towle et al., 2017; Ioannou et al., 2018). These defects may therefore offer unique insight into how different primates cope with anthropogenic-related changes to their habitat and environment.

In this study, skeletal remains of Japanese macaques (*Macaca fuscata*) from a range of locations were examined for evidence of macroscopically-visible enamel defects. All types of enamel defects were recorded, including LEH, PEH, PFEH and localized defects. A special focus was placed on individuals from the South of Yakushima Island that underwent dental development in the 1980s, since severe defects in multiple contemporary individuals were observed. Comparisons of these defects in relation to macaques from other geographic regions were made and a differential diagnosis of PFEH was undertaken, including assessment of teeth affected, potential hypo/hyper-mineralization of the remaining enamel, and tooth size/morphology comparisons. We hypothesize a severe environmental event to be responsible for PFEH defects observed.

## Materials and methods

Specimens studied originate from three Japanese islands and mainland (Yakushima, Honshu and Koshima). All specimens are curated at the Primate Research Institute (PRI) (now the Center for the Evolutionary Origins of Human Behavior), Kyoto University, Japan. All 48 individuals studied lived in the wild, with those on Koshima (Kojima) Island provisioned regularly as part of a primatological study (Watanabe, 1989; Towle et al., 2022). The data that support the findings of this study are openly available in Dryad (*doi and reference number added upon acceptance of manuscript*), which also includes the location, year of inclusion into the PRI collection and sex of each individual studied. Further details on the samples, as well as other information on tooth wear and pathologies for these populations can be found in Towle et al. (2022) and Towle and Loch (2021). This article focuses on individuals from Yakushima Island. Yakushima macaques are often considered a subspecies of Japanese macaques (*Macaca fuscata yakui*), although this is debated, especially in light of genetic evidence (Marmi et al., 2004; Fooden and Aimi, 2005). The macaques studied originate from different areas of the island (Figure 1), with a mix of males and females studied (5 males; 8 females; 8 unknown). Most individuals lived and died in the 1980s. Table 1 lists samples in more detail, including the location in which they lived.

**Figure 1.**
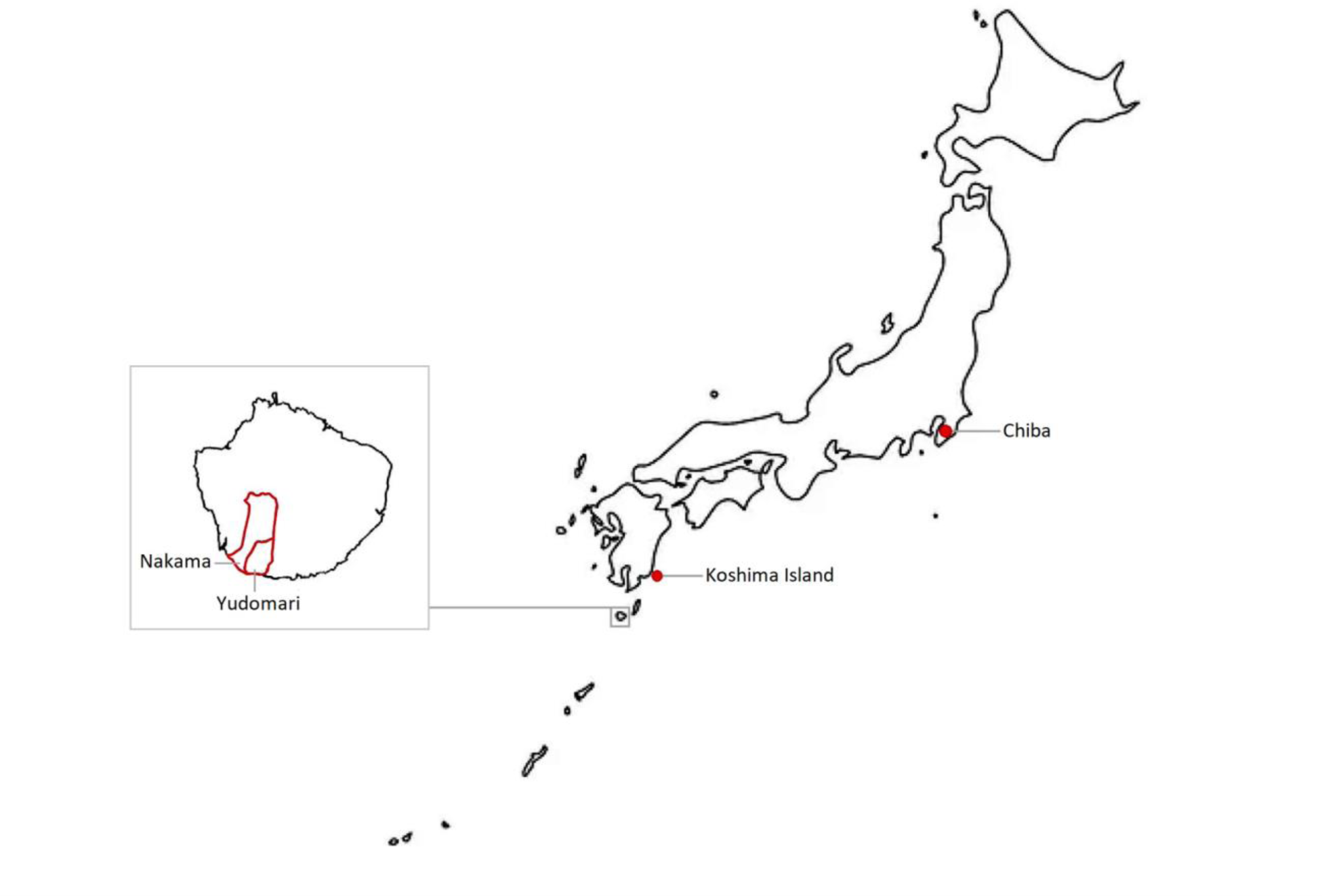
Map of Japan, with the three islands/areas from which the studied macaques lived indicated (Chiba, Koshima Island, and Yakushima Island). The two adjacent areas on Yakushima Island in which individuals with plane-form enamel hypoplasia are found are highlighted, Yudomari and Nakama.

**Table 1.** Information on the samples studied from Yakushima Island, including sample number, sex, year of death/collection/reception and locality (translated). Reception date relates to the date that the PRI study material committee received the specimen or registered it to the database.

| Sample | Sex | Reception date | Locality on Yakushima Island |
| --- | --- | --- | --- |
| 9952 | Male | 2010 | Not stated |
| 9013 | Female | 2009 | Han-yama |
| 10377 | Female | 2014 | Not stated |
| 2593 | Male | 1987 | Nakama |
| 2595 | Female | 1987 | Kusugawa |
| 2603 | Not known | 1987 | Nakama |
| 3130 | Female | 1989 | Yudomari |
| 3133 | Not known | 1989 | Yudomari |
| 2600 | Female | 1987 | Nakama |
| 2598 | Female | 1987 | Nakama |
| 3139 | Not known | Not known | Yudomari |
| 3158 | Not known | 1989 | Isso |
| 3430 | Female | 1989 | Han-yama |
| 1730 | Male | 1983 | Not stated |
| 2597 | Female | 1987 | Nakama |
| 2604 | Not known | 1987 | Nakama |
| 3124 | Male | 1989 | Yudomari |
| 3136 | Not known | 1989 | Yudomari |
| 3138 | Male | 1989 | Yudomari |
| 3435 | Not known | 1989 | Yaku-cho |
| 3434 | Not known | 1989 | Yaku-cho |

Enamel hypoplasia data was collected following Towle and Irish (2019) and summarized here. Teeth were held under a lamp and rotated allowing light to hit the surface at different angles. The smallest discernible macroscopic defect was recorded, with a hand lens used to rule out postmortem damage. Methods for recording LEH follow Goodman and Rose (1990), Guatelli-Steinberg (2003), Lukacs (1989), and Miszkiewicz (2015). Localized hypoplasia was recorded following Skinner et al. (2016). PEH was recorded if there were multiple circular/oval enamel defects on a tooth crown (Towle and Irish, 2019). If pitting was present within a LEH band, then it was recorded as LEH not PEH, but the pitting was noted. Plane-form enamel hypoplasia was recorded following Towle et al. (2017). Data are presented by tooth count rather than individual, with the number of hypoplastic teeth displayed as a percentage of the total number of observable permanent teeth.

To assess for evidence of dental tissue mineralization changes associated with PFEH, mineral concentration (MC) was calculated on two teeth of an individual with PFEH (a lower first molar without defects and a second molar with PFEH), and a tooth from an individual with no PFEH. Scans were undertaken at the PRI using a SkyScan1275 Micro-CT scanner. X-rays were generated at 100 kV, 100 µA, and 10 W, with a 1 mm Copper filter placed in the beam path. Resolution was set at 15 µm voxels, and rotation was set to 0.2-degree. Images were reconstructed using the Skyscan NRecon software (NRecon, version 1.4.4, Skyscan) with standardized settings (smoothing 3; ring artifact correction 10; beam hardening 30%). Resin-hydroxyapatite phantoms were used to calibrate grayscales and mineral densities in each specimen (Schwass et al., 2009). The calibration methods followed Schwass et al. (2009), Loch et al. (2018) and Towle et al. (2022). Average values were calculated for each individual tooth by recording MC at three locations (oval ROI: 0.15 mm diameter) across the enamel thickness (outer enamel, middle enamel and inner enamel), at four crown locations (buccal, lingual, distal, mesial). For this, a single slice was selected just above the point at which the pulp becomes visible. This standardized location was chosen because a comparisons of full crown MC is not feasible in teeth with PFEH.

The size of upper first molars was assessed in Yakushima macaques that display PFEH and those that do not, to assess if defects were associated with a reduction in crown size. Right upper first molars were selected as they do not show defects even in individuals with PFEH and therefore can be compared to other samples. Sample size was limited due to teeth with substantial wear being excluded. Each maxilla was orientated so that posterior teeth had their occlusal surface perpendicular to the optical axis of the camera. All cusps were clearly visible, and the occlusal crown area maximized. Images were uploaded into ImageJ software (Schindelin et al., 2015), and calibrated using the set scale function and through measuring the millimeter scale bar in each image. The millimeter scale was placed as close as possible to the occlusal surface of the posterior teeth, with both crown and scale in focus. Outlines were drawn around the crown base following published methodology (Wood et al., 1983; Brophy et al., 2021, Bailey et al., 2004). Mesiodistal and buccolingual diameters of the same teeth, and on the same scaled image, were measured using the straight-line tool in ImageJ and recording the maximum values for each variable (following e.g., Wood, 1991; Moggi-Cecchi et al., 2006).

## Results

Only individuals from Yakushima Island have PFEH, with 68 teeth demonstrating this type of defect. A total of 10 out of 21 individuals (48%) show these defects. Most individuals with PFEH are confirmed to have lived in the neighboring areas of Yudomari (PRI: 3130, 3133, 3139, 3124, 3136) and Nakama (PRI 2600, 2598, 2604) in the south of the island (Figure 2). Two other individuals displaying PFEH (PRI 3435 and 3434) likely came from this same area, although a precise location was not recorded. At least one more individual from Yudomari likely had PFEH defects (PRI 3138), judging by the presence of 14 dental abscesses, and an atypical wear pattern across the molars (i.e., first molars were less worn than second and third molars). This may also be the case for the other individuals from Yudomari and Nakama, with advanced wear potentially erasing evidence of previous defects. Other individuals from these two areas also display other types of enamel hypoplasia (PRI 2593, 2603, 2597). In individuals from other locations, both on Yakushima Island and Honshu and Koshima, no cases of PFEH were observed (i.e., 0/ 638 teeth for Chiba and Koshima samples combined). Other types of enamel hypoplasia were also more frequent in Yakushima Island individuals compared to Koshima Island and Chiba, including localized defects and LEH (Table 2). No individuals analyzed from any location displayed PEH.

**Figure 2.**
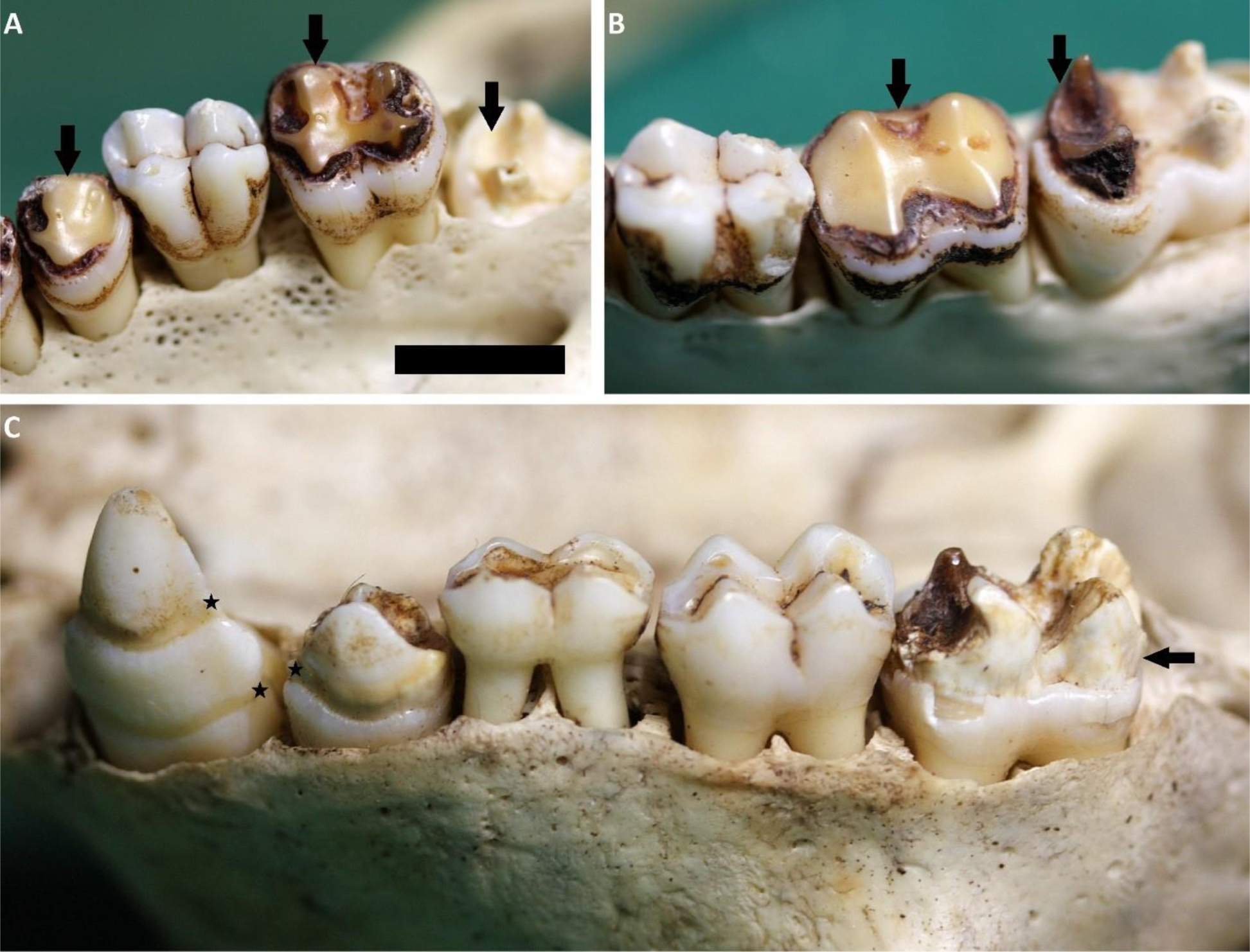
Plane-form enamel hypoplasia (PFEH) on the posterior dentition of Yakushima macaques. A) Left maxillary posterior teeth (specimen: PRI 2600), showing extensive PFEH on the fourth premolar and second and third molars (black arrows). Note the first molar appears unaffected; B) Mandibular left posterior teeth (specimen: PRI 2600), showing PFEH on both the second and third molars (black arrows). Note the first molar appears unaffected; C) Mandibular left dentition (specimen: PRI 2598), showing PFEH on the second molar (black arrow), and the canine and the third premolar show defects that are best described as PFEH, but could also be termed ‘severe LEH’ (black stars), especially the cervical most defect on the canine. Note the first molar and remaining deciduous molar are unaffected. Scale bar: 1cm.

**Table 2.**
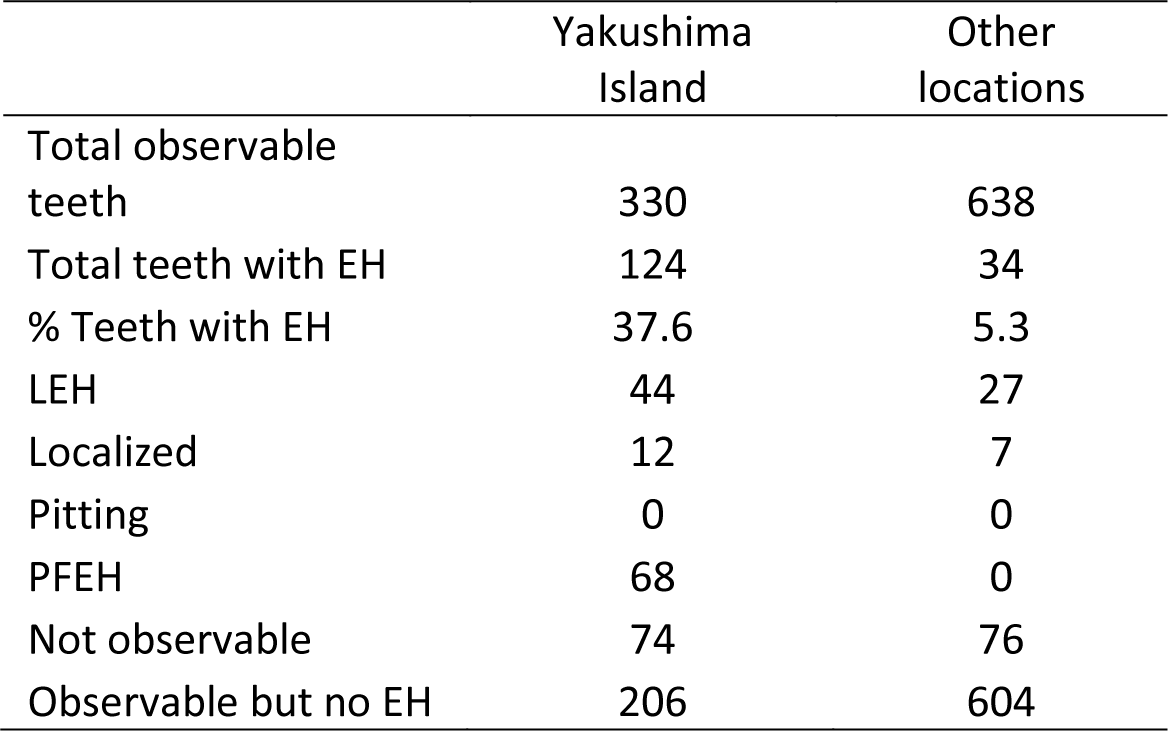
Enamel hypoplasia prevalence for all permanent teeth, split by defect type. Other locations combines figures for Chiba and Koshima samples. EH: enamel hypoplasia; LEH: linear enamel hypoplasia; Localized: localized enamel hypoplasia; Pitting: Pitting enamel hypoplasia; PFEH: plane-form enamel hypoplasia.

In individuals with PFEH, most permanent teeth are affected to varying degrees, with second molars having the highest prevalence of PFEH (Table 3). First molars were not affected by PFEH (Figure 2; Table 3). Although not studied in depth here due to small sample sizes, the low number of deciduous teeth present did not show PFEH, even if the permanent dentition did (Figure 2C). PFEH had a similar presentation across individuals and tooth types, with the most prominent examples showing absence of enamel in large areas of the crown, leaving the dentine protruding (Figures 2 and 3). In some cases, the defects did not affect the occlusal surface, with deep defects observed further down the crown (Figure 3B). On some teeth the defects appeared less severe, making it difficult to distinguish between LEH or PFEH (e.g., Figure 2C).

**Table 3.** Combined total for each tooth type (including antimeres and maxillary and mandibular teeth), for all Yakushima Island samples combined. M: molar; PM: premolar; C: canine; I: incisor. EH: enamel hypoplasia; LEH: linear enamel hypoplasia; Localized: localized enamel hypoplasia; PFEH: plane-form enamel hypoplasia.

|  | M3 | M2 | M1 | PM2 | PM1 | C | I2 | I1 |
| --- | --- | --- | --- | --- | --- | --- | --- | --- |
| Total observable teeth | 26 | 53 | 59 | 41 | 42 | 41 | 31 | 37 |
| Total teeth with EH | 9 | 24 | 5 | 15 | 19 | 14 | 14 | 24 |
| % Teeth with EH | 34.6 | 45.3 | 8.5 | 36.6 | 45.2 | 34.2 | 45.2 | 64.9 |
| LEH | 3 | 4 | 4 | 3 | 8 | 11 | 7 | 4 |
| Localized | 0 | 0 | 1 | 2 | 0 | 0 | 0 | 9 |
| PFEH | 6 | 20 | 0 | 10 | 11 | 3 | 7 | 11 |
| Not observable | 22 | 20 | 19 | 20 | 20 | 10 | 3 | 1 |
| No EH | 17 | 29 | 54 | 26 | 23 | 27 | 17 | 13 |

**Figure 3.**
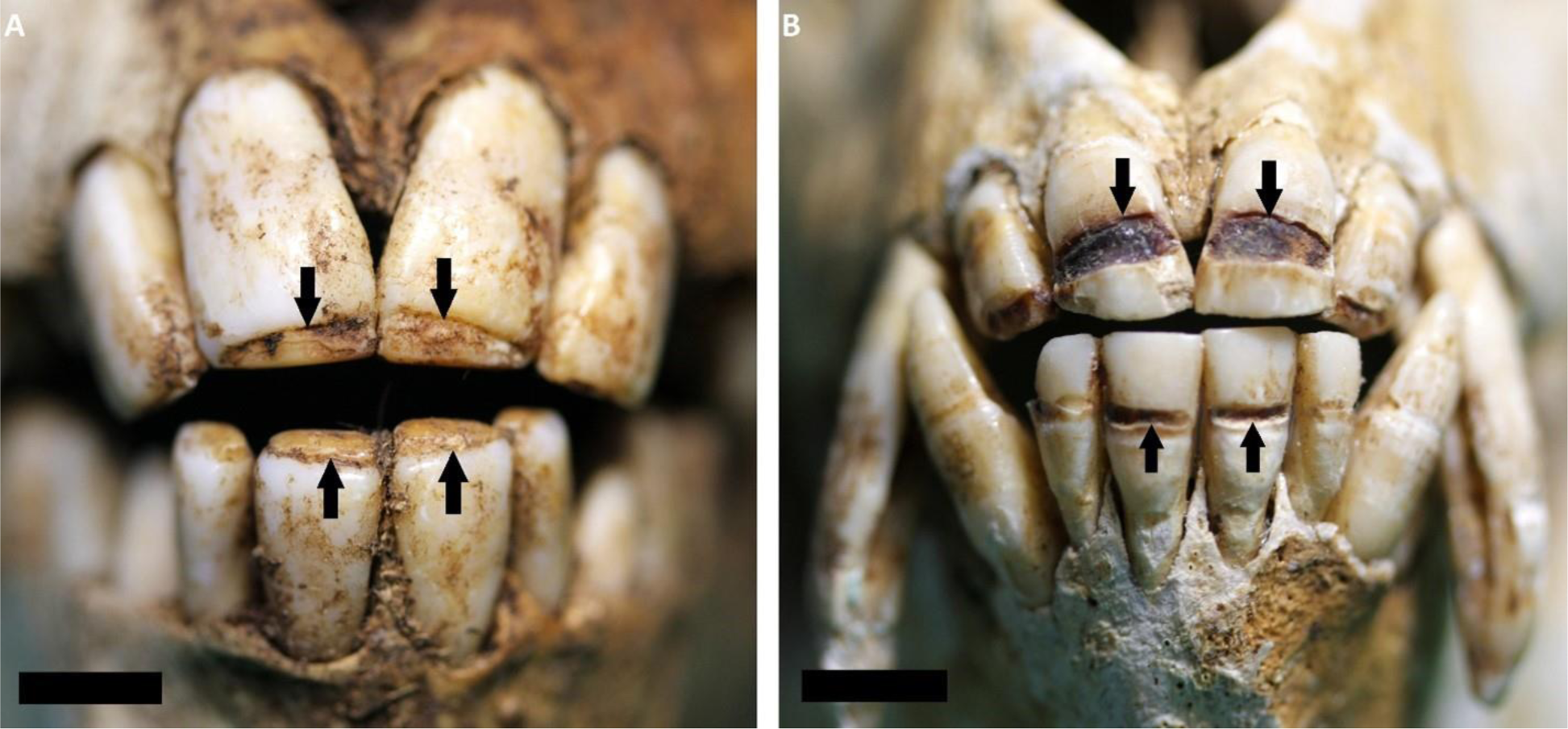
Plane-form enamel hypoplasia (PFEH) in the anterior dentition of Yakushima macaques. A) specimen PRI 2604 showing extensive PFEH defects near the incisal edge of the central incisors, leading to protruding dentine (black arrows); B) specimen PRI 3124 showing PFEH defects in the mid crown region (black arrows). Both scale bars 1cm.

There was no sign of hypo or hyper mineralization in dental tissues in the individual with PFEH analysed via Micro-CT, with MC values similar to an individual without PFEH (Table 4). There was also no evidence of extreme pathology/wear (e.g., caries or erosion) further causing demineralization of dental tissue in these areas. Individuals with no PFEH had slightly elongated upper first molars, as well as larger crown base areas (Table 5).

**Table 4.** Average mineral concentration (MC) values for enamel and dentine in three Yakushima macaque teeth: a tooth with PFEH (PRI 3434, lower right M2), without PFEH but from the same individual that has PFEH on other teeth (PRI 3434, lower right M1), and a tooth with no sign of PFEH on any tooth (PRI 1730, lower left M2).

|  | Enamel MC (g/cm<sup>3</sup>) | Dentine MC (g/cm<sup>3</sup>) |
| --- | --- | --- |
| PRI 3434 M2 (PFEH) | 2.42 ± 0.06 | 1.27 ± 0.08 |
| PRI 3434 M1 | 2.39 ± 0.05 | 1.31 ± 0.06 |
| PRI 1730 M2 | 2.46 ± 0.12 | 1.35 ± 0.03 |

**Table 5.** Mesial-distal length, buccal-lingual length and total crown base area for upper right first molars, split PFEH and no PFEH specimens

| Specimen | Sex | Mesial-distal length (mm) | Buccal-lingual length (mm) | Total crown base area (mm<sup>2</sup>) |
| --- | --- | --- | --- | --- |
| **PFEH** |  |  |  |  |
| 2600 | Female | 6.93 | 6.21 | 39.40 |
| 3139 | Not known | 7.66 | 6.41 | 39.72 |
| 2604 | Not known | 7.22 | 6.00 | 37.87 |
| 3124 | Male | 7.68 | 6.63 | 42.72 |
| 3434 | Not known | 7.30 | 6.46 | 42.89 |
| **Average** |  | **7.36** | **6.34** | **40.52** |
| **No PFEH** |  |  |  |  |
| 2595 | Female | 7.87 | 6.87 | 46.48 |
| 3158 | Not known | 7.79 | 6.97 | 45.94 |
| 3430 | Female | 7.87 | 6.46 | 44.19 |
| 1730 | Male | 8.01 | 6.54 | 44.05 |
| **Average** |  | **7.89** | **6.71** | **45.16** |

## Discussion

The results of this study suggest that the severe enamel defects observed in Japanese macaques from the southern regions of Yakushima Island were most likely caused by environmental factors and not the result of a genetic condition (i.e., amelogenesis imperfecta) or localized trauma to the developing tooth (i.e., localized enamel hypoplasia). We conclude that the defects likely relate to a severe environmental disturbance experienced by the macaques on the southern part of Yakushima Island, with permanent first molars and deciduous teeth unaffected because they form in utero and therefore would have been maternally buffered from the environmental factors that triggered the PFEH. Further, given the uniform nature of these defects across individuals, it is possible that there was a common risk-factor for the type of PFEH defect observed, such as a genetic pre-disposition brought to high frequency due to the documented bottle neck observed in the population.

There are two reasons why we conclude that these defects are not related to a genetic mutation that effects enamel generally, such as is associated with amelogenesis imperfecta (AI). First, AI typically affects all teeth, including the deciduous and permanent dentition. The PFEH observed on these macaques does not affect the first molars, and appears not to affect the deciduous dentition. Our second reason for rejecting the hypothesis of these enamel defects being a form of AI is because the quality of the remaining enamel is healthy. If hypo or hyper mineralization of enamel had contributed to the formation of these defects, or else pathology (e.g., caries) or wear (e.g., erosion) was in some way involved, then changes in the MC of the remaining dental tissue would be expected. In studies which estimated MC in hypo or hyper mineralized teeth, and others which assessed the impact of dental pathologies on dental tissue MC, a significant reduction or increase of over 0.25 g/cm^3^ has been observed compared to ‘sound’ dental tissue (e.g., Kobayashi et al., 2021; Towle et al., 2022). There is no evidence for such MC variation in the macaques analyzed here. We similarly rule out these PFEH being caused by localized enamel defects as these normally affect a certain surface or tooth and not entire tooth crowns, as is observed here (Halcrow and Tayles, 2008; Skinner et al., 2016).

This reduced tooth size could result from the shared genetic effects with body size. For example, primate tooth size is genetically correlated with body size (Hlusko et al. 2006). Yakushima macaques have several distinguishing features from other Japanese macaques, such as a smaller body size and darker coat, as well as a suite of suggested cranial morphological differences with other Japanese macaques (Kuroda, 1940; Ikeda & Watanabe, 1966; Yano et al., 2020). Genetic research suggests that these differences are likely due to a few key mutations in their genome (Lee et al., 2016; Marmi et al., 2004). Other studies have shown that the Yakushima population experienced a significant decline approximately 5000 years ago, which may have been caused by the eruption of Kikai caldera (Hayaishi and Kawamoto, 2006; Lee et al., 2016). Recent human activities including habitat destruction and culling of individuals may have further exacerbated the limited population mobility and increased the likelihood of inbreeding. This is supported by a recent study that found Yakushima macaques show extremely low genetic diversity (Ito et al., 2021). Consequently, there may be a genetic component to the observed enamel hypoplasia in the specimens evaluated in this study, possibly linked to inbreeding or pleiotropy.

The smaller tooth size of the Yakushima Japanese macaques with PFEH could also be the result of systemic stress. Other studies have shown that variation in molar size is intertwined with prenatal growth rates (Monson et al., 2022), and smaller tooth size may well be an indicator of environmental stress (DeGusta et al., 2003). The reduction in first molar size in individuals with PFEH could be related to a combination of pre- and post-natal maternal stress, with greater physiological stress experienced by individuals after birth. A study on rats found that prenatal maternal stress can significantly impact offspring growth, supporting this hypothesis (Amugongo and Hlusko, 2014). While limb bones will experience “catch-up” growth later in ontogeny, tooth crowns do not continue to grow after eruption, thereby providing a more accurate reflection of disruptions to early childhood growth (Hillson and Bond, 1997).

The high rate of PFEH could have resulted from a combination of genetic and environmental factors. Given the intense bottleneck suggested to have occurred for these macaques, and the potential for pleiotropy affects relating to recent adaptations (e.g., in coat and size phenotypes), it is feasible these groups were predisposed to developing PFEH. This would explain why such defects have not yet been described for other primates, and the similarity of defects among individuals. A somewhat similar type of group-specific enamel defect was identified in *Paranthropus robustus* (Towle and Irish, 2019) in which pitting enamel hypoplasia was commonly observed only in posterior teeth.

Logging operations intensified on Yakushima Island in the 1960s and 1970s, and many areas experienced intensification of agriculture and conifer plantations. By the 1980s, Yakushima Island macaques were regularly crop raiding in many areas, with oranges commonly consumed (Yoshihiro et al., 1998; Yamagiwa, 2008). This led to increase in macaque persecution, with trapping and culling occurring regularly (Yoshihiro et al., 1998). In Yakushima Island, the number of macaques killed or trapped between 1979 and 1988 is estimated to have been over 3,000 (Hill, 1992). Deforestation, change in forest composition and trapping did not occur uniformly across the island, with large areas of national parks retaining much of their primary forest and natural resources. The areas of focus of this study, Yudomari and Nakama, in contrast, experienced substantial changes, with native forest removed and replaced with orange and conifer plantations. It is likely that the severe PFEH observed in macaques living in these areas relate to stress due to habitat changes. These changes would have led to a reduction in diet diversity, with a likely shift to consumption of cropped plants (especially oranges). Additionally, such changes may have led to food becoming scarce at certain times of the year, potentially leading to severe malnutrition. The disturbance to social groups caused by trapping/culling could also have influenced the etiology of PFEH.

Whether habitat loss and/or environmental change are related to the occurrence of PFEH in the Yakushima Island macaques, it is difficult to piece apart the factors directly responsible for the occurrence of the enamel defects. Often, wild Japanese macaque populations will inhabit a mosaic of land types, even within small geographical ranges. Areas with primary forest, conifer plantations and naturally regenerated forests will influence and impact the behavior and diet of wild macaques (Hanya et al., 2018). This means that if enamel defects are observed, understanding the exact cause would be difficult, even if detailed foraging information was available. All of the macaques included in our study were affected by human activities in some way, including Koshima Island macaques being provisioned with food, human activities, and the introduction of non-native fauna and flora to Chiba on the mainland. Despite this, macaques in these other locations do not show severe PFEH defects or high LEH prevalence compared to other primate species in other geographic regions (Guatelli-Steinberg and Skinner, 2000; Towle and Irish, 2020). This suggests the macaques from Yudomari and Nakama must have undergone severe periods of physiological stress and severe malnutrition, or had enamel that was particularly susceptible to those stresses beyond what has been experienced by other wild primate groups.

However, other natural causes also need to be considered. Precipitation is high on Yakushima Island, and has been associated with enamel defects in other primate groups (Skinner, 2021, and reference therein). Similarly, typhoons and other severe weather events regularly affect the island, including during the period of interest (Itaka et al. 2013; Suzuki & Tsukahara 1987; Sase et al. 1998; Saito 1992). A potential example of such event affecting Yakushima macaques was described by Hanya et al. (2004), in which they report mass mortality of macaques in 1998 due to natural causes. The authors concluded the mass mortality event was likely related to severe weather (hot and dry) causing low production of fleshy fruits, leading to severe shortage of high-quality foods (Hanya et al. 2004). The authors also discussed the possibility of intertroop competition and disease, although these seem less likely to result in PFEH. Similarly, changes in home range size and group dynamics could also be linked to increased levels of physiological stress and aggression within macaque populations (Maruhashi, 1982; MacIntosh et al., 2012; Hanya et al., 2018).

Yakushima macaques offer a unique opportunity to study key questions relating to the evolution of the primate dentition, as there are noticeable size differences in their teeth and overall body size compared to other macaque populations, both on and off the island (Fooden and Aimi, 2005; this study). Previous research on primates has highlighted the crucial relationship between molar size and recent adaptation/evolution, demonstrating a complex interplay of multiple genes in dental development and accompanying changes in molar cusps with other dental traits (DeGusta et al., 2003; Koh et al., 2010; Hlusko et al., 2004). As Yakushima macaques with PFEH display smaller upper first molars and potentially altered cusp dimensions, this presents a valuable opportunity to investigate the effects of habitat change, population isolation, inbreeding, and physiological stress on dental evolution and development. Further research is necessary to distinguish normal phenotypic variation from more pathological variation related to physiological stress, especially considering the smaller size of Yakushima macaques compared to other Japanese macaque populations. This also has implications for phylogenetic interpretations, as some of the specimens identified as having PFEH in this study have been used to characterize the small size of Yakushima macaques, including smaller dental measurements (Fooden and Aimi, 2005).

Since the 1980s, mitigation of the impacts of human occupation on macaques, including plans for sustainable use of forests, have been put in place on Yakushima Island. Non-governmental organizations (NGOs) have been established to promote conservation and natural history research, and to prevent human-wildlife conflicts on the island. Further research into the health and physiological stress of these macaques through time using skeletal indicators such as enamel hypoplasia may offer further insight into the effects of human activity on these populations. Here we demonstrate that PFEH is potential tool for the study of anthropogenic impact, for which there is growing evidence of changes in diet, behavior, and social structures (e.g., Ancrenaz et al., 2014). PFEH could also be useful alongside other studies on nutrition and population health, such as previous investigations using urinalysis and fecal analysis (MacIntosh et al., 2011; MacIntosh et al., 2012), and morphological comparisons (Kamaluddin et al., 2019; Landi et al., 2021). Primatological studies have also shown how feeding behavior and ranging patterns of Yakushima macaques has changed alongside human landscape alterations, both daily and across seasons (e.g., Hanya et al., 2020). Studies on the occurrence of PFEH, with other types of research, may provide new insights into the impact of human activity on the behavior and health of Yakushima Island macaques.

PFEH is rare in wild primates and archaeological/paleontological hominin samples. This could be due to individuals being unlikely to survive such severe physiological stress events. However, this study demonstrates individuals in a wild population can survive such events. Our study suggests that substantial environmental and habitat changes can lead to severe enamel defects, highlighting a dental proxy for human-related impacts on wild primates. Further research into enamel defects may prove useful for assessing the effect of human activity in different primate groups.

## Acknowledgements

This research was supported by a Sir Thomas Kay Sidey Postdoctoral fellowship from the Faculty of Dentistry, University of Otago. This research was additionally facilitated by the European Research Council within the European Union’s Horizon Europe (ERC-2021-ADG, Tied2Teeth, project number 101054659). Views and opinions expressed are however those of the author(s) only and do not necessarily reflect those of the European Union or the European Research Council. Neither the European Union nor the granting authority can be held responsible for them. The authors thank the Study Material Committee from the Primate Research Institute (PRI), Kyoto University, for access to their collections, and T. Ito for assistance during data collection. The research was performed under the Cooperative Research Program of the PRI (2019-C-20). The authors also thank T. Ito and G. Hanya for helpful comments during the writing of this manuscript.

